# Neural correlates of conscious tactile perception: An analysis of BOLD activation patterns and graph metrics

**DOI:** 10.1101/2019.12.19.882365

**Authors:** Martin Grund, Norman Forschack, Till Nierhaus, Arno Villringer

**Author notes:** **Corresponding author:** Martin Grund, Max Planck Institute for Human Cognitive and Brain Sciences, Stephanstr. 1A, 04103 Leipzig, Germany.

## Abstract

Theories of human consciousness substantially vary in the proposed spatial extent of brain activity associated with conscious perception as well as in the assumed functional alterations within the involved brain regions. Here, we investigate which local and global changes in brain activity accompany conscious somatosensory perception following electrical finger nerve stimulation, and whether there are whole-brain functional network alterations by means of graph metrics. Thirty-eight healthy participants performed a somatosensory detection task and reported their decision confidence during fMRI. For conscious tactile perception in contrast to undetected near-threshold trials (misses), we observed increased BOLD activity in the precuneus, the intraparietal sulcus, the insula, the nucleus accumbens, the inferior frontal gyrus and the contralateral secondary somatosensory cortex. For misses compared to correct rejections, bilateral secondary somatosensory cortices, supplementary motor cortex and insula showed greater activations. The analysis of whole-brain functional network topology for hits, misses and correct rejections, did not result in any significant differences in modularity, participation, clustering or path length, which was supported by Bayes factor statistics. In conclusion, for conscious somatosensory perception, our results are consistent with an involvement of (probably) domain-general brain areas (precuneus, insula, inferior frontal gyrus) in addition to somatosensory regions; our data do not support the notion of specific changes in graph metrics associated with conscious experience. For the employed somatosensory submodality of fine electrical current stimulation, this speaks for a global broadcasting of sensory content across the brain without substantial reconfiguration of the whole-brain functional network resulting in an integrative conscious experience.

## Introduction

In the debate on the neural correlates of consciousness, several crucial issues are still not resolved. First, regarding the involved brain regions, some studies assume only areas related to the particular perceptual modality to be necessary (Auksztulewicz et al., 2012; Schröder et al., 2019), others emphasize the role of a parietal hot zone (Boly et al., 2017; Koch et al., 2016) whereas some theories - most notably the global workspace theory (Baars, 1988; Dehaene et al., 2006; Mashour et al., 2020) - assume conscious experience to depend on the involvement of widespread particularly fronto-parietal brain regions (Naghavi and Nyberg, 2005; Rees et al., 2002). Second, regarding the neurophysiological processes occurring *within* the involved brain regions, recent studies are suggesting specific alterations in their connectivity which can be identified by effects of transcranial stimulation (Casali et al., 2013) or using graph-theoretical metrics on fMRI data (Godwin et al., 2015; Sadaghiani et al., 2015).

For the somatosensory domain, previous fMRI activation studies have suggested the ipsilateral and contralateral secondary somatosensory cortices as the most promising candidates for conscious tactile perception (Moore et al., 2013; Schröder et al., 2019). Furthermore, recurrent interaction of S2 with S1 may play an important role for tactile detection (Auksztulewicz et al., 2012). Research focusing on tactile illusions has shown that S1 is activated somatotopically in correspondence with the illusory percept and body ownership (Blankenburg et al., 2006; Martuzzi et al., 2015). In this context, the temporal parietal junction (TPJ) plays a major role for bodily self-consciousness (De Ridder et al., 2007; Ionta et al., 2014). The insula has been consistently reported to be associated with conscious tactile perception (Moore et al., 2013) and described as a central hub for interoception (Ronchi et al., 2015) and self-identification (Park and Blanke, 2019). Interestingly, in another recent study, the insula together with anterior cingulate cortex coded for uncertainty across stimulation intensities (Schröder et al., 2019). In the same study, frontal and parietal activations in tactile detection paradigms have been interpreted as serving the task (e.g., reporting a percept) but not the conscious sensory experience (Schröder et al., 2019). Yet, the above-mentioned ideas of a “global workspace” involving mainly fronto-parietal activity (Dehaene et al., 2006), or of a posterior cortical hot zone integrating sensory cortices (Koch et al., 2016) are conceived to be domain-general thus also applying to the tactile consciousness.

While most of the above-mentioned studies relied on the analysis of BOLD activation patterns, the question, whether functional connectivity changes or not, can be assessed using graph metrics (Bassett and Sporns, 2017). For this purpose, cortical and subcortical regions of interest (ROIs) are defined as nodes and their temporal relationships as edges (i.e., their connection; Bullmore and Bassett, 2011). The resulting network topologies are assessed with graph-theoretical measures such as modularity and clustering coefficient and compared between aware and unaware target trials (Godwin et al., 2015; Sadaghiani et al., 2015; Weisz et al., 2014). *Modularity* captures the global organization of nodes in subnetworks, whereas the *clustering coefficient* indicates whether a node’s neighbors are also connected, thus forming local clusters. Measures of integration (e.g., *characteristic path length*) describe the general connectivity between all nodes, whereas measures of centrality (e.g., *participation coefficient*) reveal important nodes in the network. In this framework, visual awareness has recently been suggested to be accompanied by a decreased modularity and increased participation coefficient of the post-stimulus network topology in fMRI (Godwin et al., 2015). Importantly, these topologies had explanatory power beyond local BOLD amplitudes and baseline functional connectivity (Godwin et al., 2015). Globally, this indicates a lower segregation of nodes into distinct networks and locally a higher centrality of all nodes. A more integrated state accompanying stimulus awareness (Godwin et al., 2015) is supposed to facilitate broadcasting of sensory information to other brain areas (Dehaene et al., 2006; Dehaene and Changeux, 2011). These widespread changes in functional connectivity have been interpreted as evidence supporting global models of awareness, e.g., the global workspace theory (Dehaene et al., 2006; Dehaene and Changeux, 2011). Whether these changes in graph metrics generalize to other sensory modalities is not yet answered.

In the present study, building on our previous experience in studies on neural processes underlying conscious and unconscious somatosensory processing (Blankenburg et al., 2003; Forschack et al., 2020; 2017; Nierhaus et al., 2015; Schubert et al., 2006), we used a “classical” fMRI detection paradigm in which aware and unaware trials of physically identical near-threshold stimuli are contrasted (Aru et al., 2012; Baars, 1988). Notably, our fMRI paradigm included a nine-second pause between stimulation and report, which made the assessment of functional network topologies with graph metrics possible. Therefore, our study design allowed to assess BOLD activity patterns and graph metrics independently, as well as to relate the two measures directly.

Two other features of our paradigm are important: (i) a confidence rating was included for each trial allowing for an analysis of confident decisions only and (ii) the paradigm included 25% catch trials. By comparing the contrast of undetected stimuli to correctly rejected catch trials, neural processes associated with non-conscious stimulus processing of near-threshold stimuli can be assessed. In a previous study on subthreshold stimuli, we had shown that they were associated with a deactivation of somatosensory brain regions (Blankenburg et al., 2003); however, it is not clear whether this is also true for stronger stimuli near the detection threshold.

Thus, our study aimed to address the following main questions:

- Do BOLD activation patterns following somatosensory near-threshold stimuli match the predictions of major consciousness theories, i.e., does the contrast detected/undetected lead to increased activity only in somatosensory areas (Schröder et al., 2019), in a fronto-parietal network (global workspace theory), or in a more restricted temporo-parietal-occipital hot zone (Koch et al., 2016)?
- Do graph metrics change with the conscious experience of somatosensory stimuli as shown for the visual system by Godwin et al. (2015), and do the affected areas match activated brain areas?
- Which neural changes are associated with non-perceived, but near-threshold somatosensory stimuli?

## Materials & Methods

### Participants

Thirty-eight healthy humans (19 women; mean age = 27.3, age range: 23-36) participated in the study. They had normal or corrected-to-normal vision and were right-handed (mean laterality index = 85, range: 60-100; Oldfield, 1971).

### Ethics statement

All participants provided informed consent (including no contra-indication for MRI), and all experimental procedures were approved by the ethics commission at the medical faculty of the University of Leipzig.

### Experimental design and statistical analysis

The experimental design of the tactile detection task had the intention to generate different sensory experiences for physically identical stimulus presentations. Brain activity accompanying these sensory experiences was sampled with BOLD fMRI (see fMRI data acquisition for details). The tactile stimulation was applied as a single electrical pulse to the left index finger. The stimulation intensity was set to the individual sensory threshold, before each of the four acquisition blocks, such that participants reported a stimulus detection (“hit”) in about 50% of the trials. One hundred near-threshold trials were intermingled with 20 clearly perceivable, supra-threshold trials and 40 catch trials without stimulation as control conditions. Participants had to report their perception (yes/no) and decision confidence (see “behavioral paradigm” for details). This led to three within-participant factors of interest: (a) rejected catch trials without stimulation (correct rejections), (b) non-perceived near-threshold trials (misses), and (c) perceived near-threshold trials (hits). We did not include false alarms (reported “yes” in catch trials without stimulation) due to the low false alarm rate (mean FAR = 3.3%, *SD* = 6.0%). 17 of 31 participants reported zero false alarms.

We compared the graph metrics between hits, misses and correct rejections across participants with the Wilcoxon’s signed-rank test (see Graph-theoretical analysis for details). For each graph metric, the *p*-values of the 24 paired Wilcoxon’s signed-rank tests were corrected for multiple comparisons with a false discovery rate (FDR) of 5% (Benjamini and Hochberg, 1995). The BOLD response amplitudes were modeled for the three (detection related) within-participant factors and compared them with a mixed-effects meta-analysis (3dMEMA; Chen et al., 2012). We controlled for multiple comparisons with family-wise error correction (see fMRI contrast analysis for details).

### Data and code availability

The code to run the experiment, the behavioral data, and the code to analyze the behavioral and MRI data are available at http://github.com/grundm/graphCA. Due to a lack of consent of the participants, structural and functional MRI data cannot be shared publicly, and can only be made available upon reasonable request if data privacy can be guaranteed according to the rules of the European General Data Protection Regulation (EU GDPR). The respective research group has to sign a data use agreement to follow these rules. This statement is in line with our institute’s policies and requirements by our funding bodies.

### Behavioral paradigm

Participants had both to report the perception (yes/no) of electrical pulses applied to their left index finger and rate their confidence about their decision. Single square-wave pulses (0.2 ms) were generated with a constant current stimulator (DS5; Digitimer, United Kingdom) at individually assessed intensities near (mean intensity = 1.85 mA, range: 1.01-3.48 mA) and supra (mean intensity = 2.18 mA, range: 1.19-3.89 mA) perceptual threshold reflecting 50% and 100% detection rate. Additionally, 25% of all trials were catch trials without stimulation.

Each trial (21 s) started with a fixation cross (1 s), followed by a cue (1 s) indicating an electrical pulse was soon to follow (Figure 1). The stimulation onset was always 100 ms before cue offset in order to temporally align the stimulation with the detection decisions. For aware trials participants’ detection decisions presumably occur the instant the stimulation is noticed. However, for unaware trials they can only conclude there was no stimulus at cue offset. If the stimulus onsets had been pseudo-randomized across the cue window, the yes-decision would have occurred on average half of the cue window earlier than the no-decision. The actual reporting of the decision was delayed by 9 s to allow a movement-free time window for analyses. Participants had 1.5 s to report if they felt the stimulus or not by pressing the corresponding button for yes or no. Subsequently they had another 1.5 s to report their confidence about the yes/no-decision on a scale from 1 (very unconfident) to 4 (very confident). Any remaining time in the confidence rating window, following the rating, was added to a 7 s fixation cross creating an inter-trial interval of at least 7 s. Participants were instructed to place their right four fingers on a four-button box. The second and third buttons were controlled by the right middle finger and the ring finger to report the decision for yes or no. The outer buttons were controlled by the index finger and the little finger additionally to report the confidence decision on the full four-point scale. All button-response mappings were counterbalanced across participants. Hence depending on the mapping, the middle finger or the ring finger indicated “yes”, and the four-point confidence scale started with “very confident” or “very unconfident” at the index finger.

**Figure 1.**
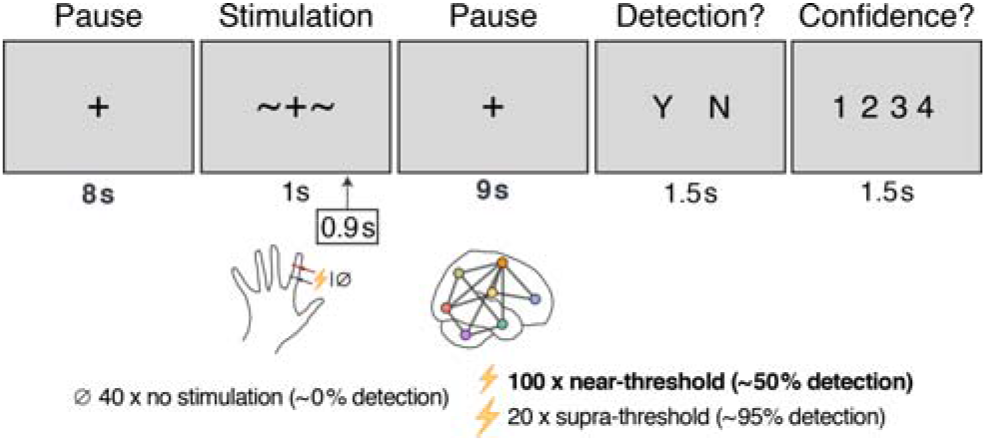
Experimental paradigm visualized across one trial (21 s). The tiles represent the participant’s visual display and the times given below indicate the presentation duration. In total, each participant completed 160 trials across 4 blocks, including 100 near-threshold trials. Electrical nerve stimulation was applied to the left index finger 0.9 s after cue onset (~+~) to temporally align yes/no-decisions, which presumably had to be made at cue offset. Participants only reported their target detection decision (yes/no) following a long pause of 9 s in order to model the brain’s post-stimulus functional network without a button press-related signal. The detection decision was followed by a confidence rating on a scale from 1 (very unconfident) to 4 (very confident). Every 0.75 s, a full MRI brain volume (BOLD) was acquired with a 3-mm isotropic resolution.

Each block had in total 40 trials and lasted 14 min: 10 trials without stimulation, 25 with near-threshold intensity, and 5 with supra-threshold intensity, delivered in pseudo-randomized order. Before each block, individual thresholds were assessed with an up-and-down method followed by the psi method from the Palamedes Toolbox (Kingdom and Prins, 2009). The threshold procedure followed that of the actual experimental trials but excluded the long pause and confidence response. Participants performed 4 blocks sequentially (circa 80 min). The experimental procedure was controlled by custom MATLAB scripts (The MathWorks Inc., Natick, Massachusetts) using Psychophysics Toolbox (Kleiner et al., 2007).

### fMRI data acquisition

While participants performed the task, we acquired whole-brain BOLD contrast images with a 32-channel head coil on a Siemens MAGNETOM Prisma 3 Tesla scanner. For sub-second, whole-brain images, we used a Multi-Band (MB) echo-planar imaging (EPI) sequence (Moeller et al., 2010; Setsompop et al., 2012) with an MB acceleration factor of 3 (TR = 750 ms, TE = 25 ms, flip angle = 55°, receiver bandwidth = 1815 Hz/px, partial Fourier = 7/8). No GRAPPA acceleration was applied (iPAT factor = 1). In each of the 4 blocks we acquired 1120 brain volumes (14 min), each consisting of 36 axial slices with a field of view of 192 x 192 mm² (64 x 64 voxel) and a 0.5-mm gap resulting in 3-mm isotropic voxels.

For magnetic distortion correction of the EPI scans, B0 images were obtained from double-echo GRE images (TR = 750 ms, TE_1_ = 4.92 ms, ΔTE = 2.46 ms, echo spacing = 0.66 ms, flip angle = 45°), with the same voxel geometry as used for the EPI scans. The receiver bandwidth was 822 Hz/pixel.

For normalizing the EPI scans and extracting nuisance regressors of core white matter voxels and ventricles, we used previously acquired T1-sensitive brain images of the participants with a 32-channel head coil or for two participants with a 20-channel head/neck coil on 3-Tesla Siemens MAGNETOM Prisma, Skyra, TrioTim or Verio scanner. The MPRAGE sequence covered the whole brain (176 × 240 × 256 m^3^) with an isotropic voxel resolution of 1 mm and slightly varied regarding the echo time and the receiver bandwidth across participants (TR = 2.3 s, TE = [2.01 (2), 2.96 (9), 2.98 (19), 4.21 (5)] ms, inversion time TI = 900 ms, flip angle = 9°, bandwidth = [238 (10), 240 (16), 241 (9)] Hz/px). For two participants, the sequence parameters were more different (TR = 1.3 s, TE = 3.5 ms, inversion time = 650 ms, flip angle = 8°/10°, bandwidth = 190 Hz/px).

### Behavioral data analysis

The behavioral data was analyzed with R 3.6.0 in RStudio 1.2.1335. Data by four participants was incomplete due to technical issues and failed data acquisition. The blocks of the remaining 34 participants were evaluated for successful near-threshold assessments if at least 4 null and 17 near-threshold trails with a yes/no and confidence response were recorded. This meant that only blocks with a hit rate at least 5 percentage points larger than the false alarm rate and participants with an average hit rate of 20-80% were further processed. This resulted in 31 participants with on average 89 near-threshold trials (range: 66-100). The distribution of mean detection rates is visualized in Figure 2a. For the confidence ratings, we calculated conditional probabilities for each confidence rating given a stimulus-response condition: correct rejection, near-threshold miss, near-threshold hit, and supra-threshold hit (Figure 2b). The conditional probabilities were compared with paired *t-*tests between neighboring conditions (correct rejection vs. near-miss, near-miss vs. near-hit, near-hit vs. supra-hit). The twelve *p*-values were FDR-corrected with a false discovery rate of 5% (Benjamini and Hochberg, 1995) and visualized with the means in Figure 2b.

**Figure 2.**
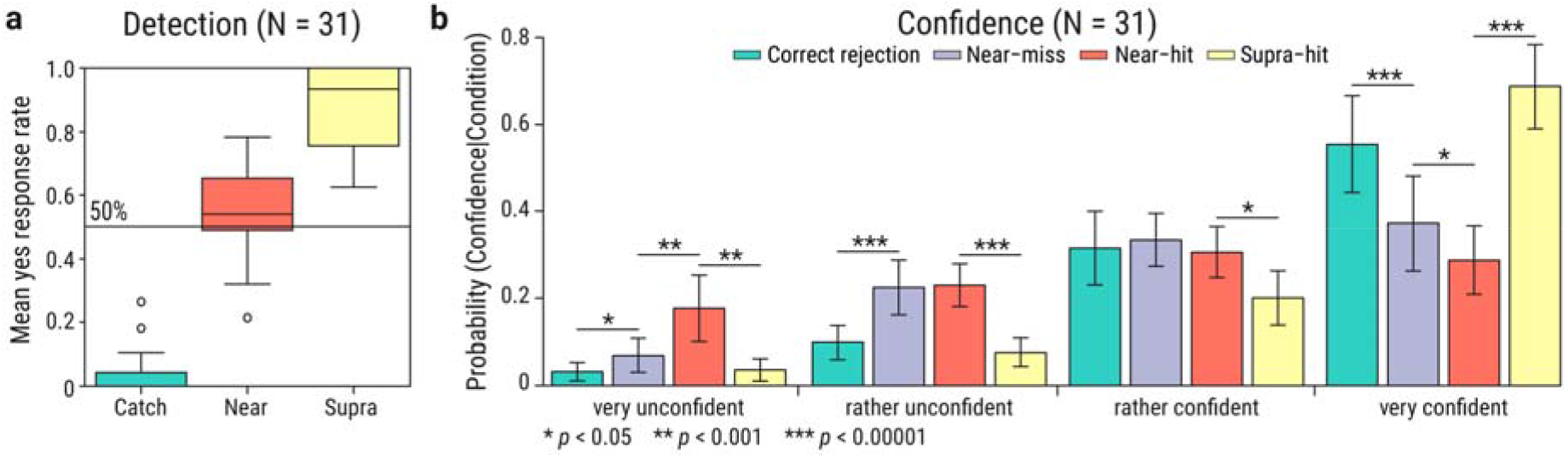
Mean detection rate and decision confidence across participants. (**a**) Detection rates for each trial condition: without stimulation (catch trials) and with near- and supra-threshold stimulation. The central line indicates the median in each box. The whiskers indicate 1.5 times the interquartile range or the maximum value if smaller. Circles indicate values beyond this whisker range. (**b**) Mean conditional probabilities for each confidence rating given a stimulus-response condition: correct rejection (green), near-threshold misses (purple), near-threshold hits (red), and supra-threshold hits (yellow). Error bars indicate within-participants 95% confidence intervals (Morey, 2008). Horizontal lines indicate significant paired *t-*tests with FDR-corrected *p*-values between neighboring conditions (Benjamini and Hochberg, 1995).

### fMRI preprocessing

Each EPI block was preprocessed with custom bash scripts using AFNI 18.2.17, FSL 5.0.11, and FreeSurfer 6.0.0 (Cox, 1996; Fischl, 2012; Smith et al., 2004). The code is available on http://github.com/grundm/graphCA. After removing the initial 10 volumes, the time series were despiked and corrected for slice timing. Subsequently, the volumes were corrected for motion and distortion using field maps acquired at the beginning of the experiment. We applied a non-linear normalization to MNI space (AFNI 3dQwarp). Next to the realignment to correct for motion, we calculated the euclidean norm (enorm) to censor volumes with large motion for the functional connectivity and BOLD contrast analyses. Volumes were ignored when they exceeded motion > 0.3 mm (enorm = sqrt(sum squares) of motion parameters; AFNI 1d_tool.py -censor_motion). Compared to the framewise displacement (FD = sum(abs) of motion parameters; Power et al., 2012), the euclidean norm has the advantage to represent appropriately large motion, e.g., the six parameters “6 0 0 0 0 0” and “1 1 1 1 1 1” would be the same for FD (FD = 6) in contrast to a enorm of 6 and 2.45, respectively. Modeling the functional connectivity and the BOLD contrasts was done with AFNI 19.1.05.

### fMRI whole-brain contrast analysis

For the BOLD contrast analysis, the data was additionally smoothed with a 7-mm FWHM kernel and scaled to a mean of 100 and maximum of 200. In the final step, we calculated a nuisance regression to control for (a) motion with Friston’s 24-parameter model (Friston et al., 1996), (b) signal outliers and their derivatives, (c) each 3 first principal components of core voxels in ventricle and white matter masks separately, and (d) a constant and trends up to polynomial of degree six (~high-pass filter > 0.0046 Hz) separately for each block.

We calculated an individual general linear model (GLM) for each participant with AFNI 3dREMLfit that combined all blocks and modeled the BOLD response as a gamma function for the following conditions: correct rejections, near-threshold misses, and near-threshold hits. A second model included confident correct rejections, confident misses, and confident hits (pooled confidence ratings of 3 and 4). Furthermore, two BOLD response regressors for the button presses of the yes/no-decision and the confidence rating were included. The regressors of the nuisance regression served as baseline regressors (AFNI 3dDeconvolve -ortvec).

The estimated regression coefficients for the aware and unaware condition were tested against each other with a mixed-effects meta-analysis (3dMEMA; Chen et al., 2012). This approach accounts for within-participant variability by using the corresponding *t*-statistics of the regression coefficients from each participant. Additionally, the detection rate was used as a covariate to account for the interindividual variance. The resulting volumes with *t*-values were corrected for multiple comparisons by thresholding voxels at *p*_voxel_ < 0.0005 and the resulting clusters at *k* voxels (*p*_cluster_ = 0.05). The cluster size threshold *k* was derived for each contrast separately based on 10,000 simulations without a built-in math model for the spatial autocorrelation function as recommended by AFNI (for details see 3dttest++ with Clustsim option and Cox et al. (2017) as response to Eklund et al. (2016)). The rendered brain images were created with MRIcron (Rorden and Brett, 2000).

### fMRI contrast analysis in primary somatosensory cortex

Furthermore, we wanted to evaluate the BOLD signal for the near-threshold stimulation in the primary somatosensory cortex. Unlike the fMRI contrast analysis for the whole brain described above, the BOLD data was not smoothed, scaled or part of a nuisance regression. We modeled the BOLD response for all near-threshold trials and trials without stimulation (independent of the yes/no-responses). The GLMs also included one regressor for each button press. The baseline regressors were limited to (a) Friston’s 24-parameter model, (b) signal outliers and their derivatives, and (c) a constant and a linear trend separately for each block (polynomial of degree one). The estimated regression coefficients for the trials with and without near-threshold stimulation were compared with a mixed-effects meta-analysis (3dMEMA; Chen et al., 2012) that included the detection rate as a covariate. Additionally to reporting the contrast “stimulus present > absent” for the whole-brain, this analysis was limited to the right primary somatosensory cortex (Area 3b and Area 1) as defined by a multi-modal parcellation based brain atlas (Glasser et al., 2016).

### Functional connectivity analysis

For estimating the context-dependent functional connectivity between regions of interest (ROI), we used the generalized psychophysiological interaction (gPPI; McLaren et al., 2012) without the deconvolution step, as implemented in FSL (O’Reilly et al., 2012). The deconvolution algorithm tries to estimate the underlying neural activity to match it temporally with the psychological context (Cisler et al., 2014; Gitelman et al., 2003; McLaren et al., 2012). However, it cannot be determined if this estimate is correct (Cole et al., 2013; O’Reilly et al., 2012). Furthermore, also Godwin et al. (2015) repeated their analysis without the deconvolution step. Hence, we followed the FSL implementation and convolved the psychological variable with a fixed-shaped HRF to temporally align it with the measured BOLD signal (O’Reilly et al., 2012). The gPPI model included (a) regressors for the BOLD response function for each condition, (b) a regressor for the baseline functional connectivity of a seed region of interest (ROI), and (c) regressors for the context-dependent functional connectivity of the ROI for each condition (psychophysiological interaction). For (b), the seed ROI average time series was extracted to be used as a regressor. For (c), this baseline regressor was masked for each condition separately to generate conditional interaction regressors. The mask for each condition was equivalent to the regressor that modeled the BOLD response for the corresponding condition, hence weighting the seed time series in the post-stimulus phase with the hemodynamic response. The interaction regressors for each condition allowed the estimation of (c) the context-dependent functional connectivity by accounting for (a) the BOLD response and (b) the baseline functional connectivity (Figure 5b). Additionally, the gPPI included baseline regressors: (a) Friston’s 24-parameter model for motion, (b) signal outliers and their derivatives, and (c) a constant and a linear trend separately for each block (polynomial of degree one).

The gPPI was calculated with AFNI 3dREMLfit for a whole-brain network of 264 nodes based on a resting-state functional connectivity atlas (Power et al., 2011). The nodes were defined as 4-mm radius, spherical ROIs at the atlas’ MNI coordinates. The BOLD response model was a gamma function. AFNI 3dREMLfit has the advantage of allowing for serial correlations by estimating the temporal autocorrelation structure for each voxel separately.

For each node’s gPPI, the coefficients of the context-dependent functional connectivity regressors were extracted from all other nodes separately by averaging across all voxels constituting the particular node. Subsequently, the beta values were combined in a symmetric connectivity matrix for each participant and each condition. As Godwin et al. (2015), we did not assume directionality and averaged the absolute values of reciprocal connections. Subsequently, the connectivity matrices were thresholded proportionally for the strongest connections and rescaled to the range [0,1] by dividing all values by the maximum value. The figures showing nodes and edges on a glass brain (Figure 5a,e) were created with BrainNet Viewer 1.6 (Xia et al., 2013).

After running the functional connectivity analysis as Godwin et al. (2015) for only confident trials, we repeated the analysis for all trials independent of their confidence response. Furthermore, we extended the preprocessing to include 7-mm smoothing, scaling and a nuisance regression and redid the analysis for both trial selections (confident only and all). For this analysis, the baseline regressors were (a) Friston’s 24-parameter model, (b) signal outliers and their derivatives, (c) each three first principal components of core voxels in ventricle and white matter masks separately, and (d) separately for each block a constant and trends up to polynomial of degree six (~high-pass filter > 0.0046 Hz).

### Graph-theoretical analysis

The context-dependent connectivity matrices were further processed with the Brain Connectivity Toolbox (BCT Version 2017-15-01; Rubinov and Sporns, 2010) to describe their network topologies. Across proportional thresholds (5-40%) graph metrics were calculated and normalized with the average graph metrics of 100 random networks with the same degree distribution (see BCT function randmio_und.m on https://sites.google.com/site/bctnet/Home/functions). In order to compare our results with the report for visual awareness (Godwin et al., 2015), we chose the same metrics for (a) segregation, (b) integration, and (c) centrality: (a) weighted undirected modularity (BCT function modulartiy_und.m; Newman, 2004) and weighted undirected clustering coefficient averaged over all nodes (BCT function clustering_coef_wu.m; Onnela et al., 2005), (b) weighted characteristic path length (BCT function charpath.m), and (c) weighted participation coefficient averaged over all nodes (BCT function participation_coef.m; Guimerà and Nunes Amaral, 2005). The participants’ graph metrics were compared between each condition with the Wilcoxon’s signed-rank test because the distributions of the graph metrics are unknown. The resulting 24 *p*-values for each graph metric (8 network threshold times 3 comparisons: hit vs. miss, hit vs. correct rejection, and miss vs. correct rejection) were FDR-corrected with a false discovery rate of 5% (Benjamini and Hochberg, 1995). Furthermore, we calculated the Bayes factors based on *t*-tests with a JZS prior (*r* = √2/2) to assess the evidence for the null hypothesis (Rouder et al., 2012).

## Results

### Behavioral data

Participants (*N* = 31) detected on average 55% of the near-threshold pulses (*SD* = 13%), 88% of the supra-threshold pulses (*SD* = 12%), and correctly rejected 97% of the catch trials without stimulation (*SD* = 6.0%; Figure 2a). Participants reported on average to be “rather confident” or “very confident” for 87% of the correct rejections (*SD* = 13%), 70% of the near-threshold misses (*SD* = 23%), 59% of the near-threshold hits (*SD* = 27%) and 89% of the supra-threshold hits (*SD* = 13%). Participants reported significantly more often “very confident” for near-threshold misses (*M* = 37.2%) than hits (*M* = 28.7%, FDR-corrected *p* = 0.037) and less often “very unconfident” for misses (*M* = 6.9%) than hits (*M* = 17.7%, FDR-corrected *p* = 0.023; Figure 2b). The conditional probabilities for “rather unconfident” and “rather confident” did not differ between near-threshold hits and misses. Near-threshold misses and correct rejections differed in their conditional probabilities for “very unconfident”, “rather unconfident” and “very confident” (Figure 2b) indicating higher confidence for correct rejections. Also, participants were on average more confident for supra-threshold hits than near-threshold hits. Additionally, we assessed the stability of near-threshold detection and false alarms across the experiment. We used linear mixed-effects models with maximum likelihood estimation (lmer function in R) to investigate the effect of block on near-threshold hit rate (near_yes) and false alarm rate (null_yes). Model comparison of the model “near_yes ~ block + (1|ID)” with the null model “near_yes ~ 1 + (1|ID)” resulted in no significant difference (χ*²* = 1.40, *p* = 0.24), indicating no effect of block on near-threshold hit rate. Also, for false alarms, the linear mixed-effects model “null_yes ~ block + (1|ID)” was not significantly different compared to the null model “null_yes ~ 1 + (1|ID)” (χ*²* = 1.41, *p* = 0.23). Therefore, we conclude that the behavioral performance is not affected by acclimatization or mental fatigue.

### BOLD amplitude contrasts

First, we modeled the BOLD contrast between hits and misses (Figure 3a-c), as well as misses and correct rejections independent of the confidence rating (Figure 3d-f). Second, we compared only confident hits and misses (Figure 4a-c), as well as confident misses and correct rejections (Figure 4d-f). Third, we modeled the contrast between near-threshold stimuli and trials without stimulation for the whole brain (Figure 5) and the primary somatosensory cortex only (Figure 6). The preprocessing for this contrast excluded smoothing, scaling or nuisance regression. For all group-level comparisons, we used the detection rate as a covariate to account for the interindividual variance (Figure 2a).

**Figure 3.**
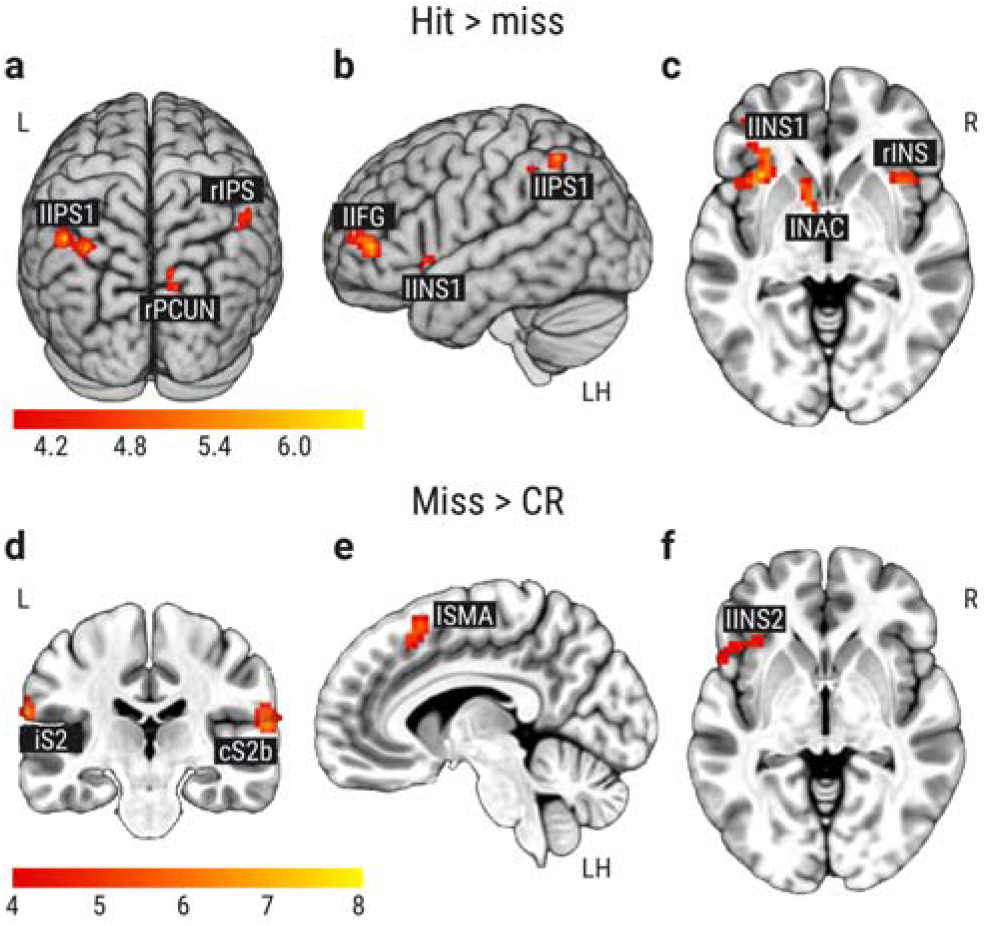
BOLD amplitude contrasts for awareness and stimulation effect. (**a-c**) Contrast between near-threshold hits and misses with focus on (**a**) the right precuneus (rPCUN) and the left and right intraparietal sulcus (lIPS1, rIPS1), (**b**) the left inferior frontal gyrus (lIFG) and (**c**) the left nucleus accumbens (lNAC) and the left and right anterior insula (lINS1, rINS; *z* = −3). Correction for multiple comparison with *t*_voxel_(30) ≥ 3.92, *p*_voxel_ ≤ 0.0005 and cluster size *k* ≥ 31 (*p*_cluster_ ≤ 0.05). (**d-f**) Contrast between near-threshold misses and correct rejections (CR) of trials without stimulation. (**d**) Coronal view (*y* = −29) with the contralateral and ipsilateral secondary somatosensory cortices (cS2, iS2). (**e**) Sagittal view (*x* = −7) on the supplementary motor area (SMA). (**f**) Axial view (*z* = −3) on the left anterior insula (INS). Correction for multiple comparison with *t*_voxel_(30) ≥ 3.92, *p*_voxel_ ≤ 0.0005 and cluster size *k* ≥ 27 (*p*_cluster_ ≤ 0.05). Left (L), right (R), and the left hemisphere (LH) are indicated.

**Figure 4.**
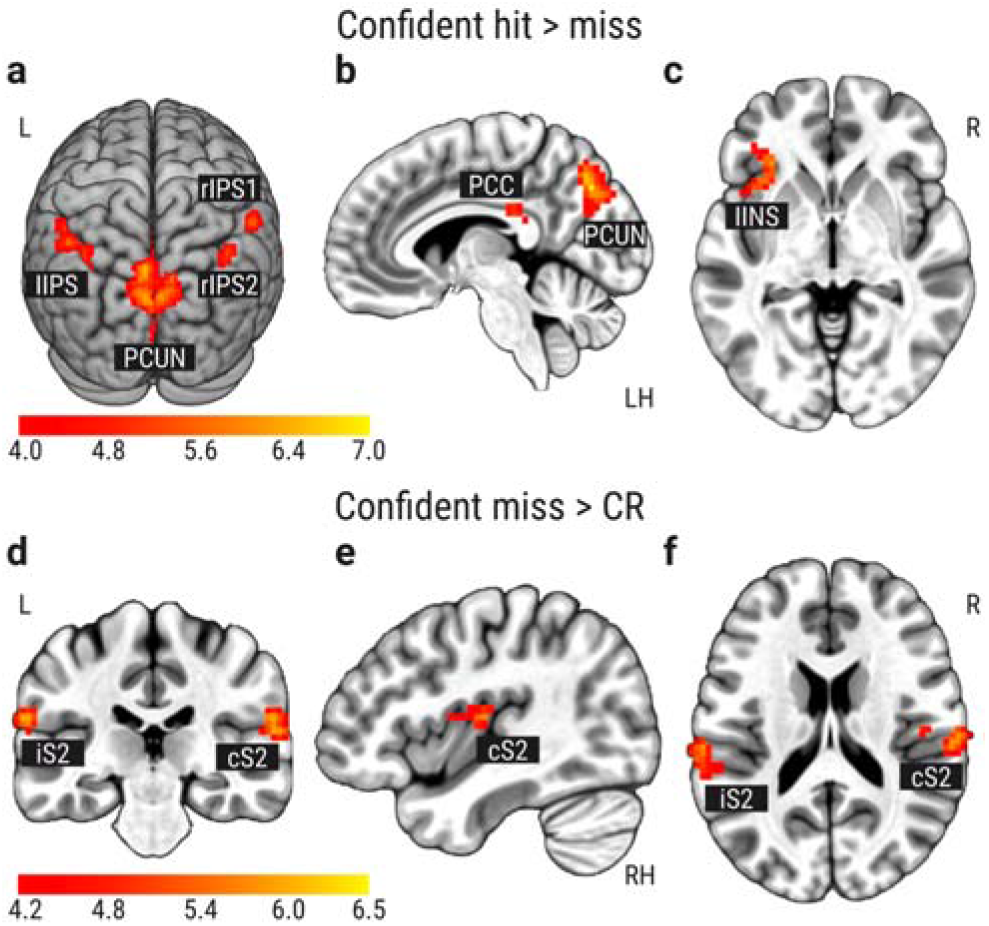
BOLD amplitude contrasts for only confident trials. Correction for multiple comparison with *t*_voxel_(30) ≥ 3.92, *p*_voxel_ ≤ 0.0005 and cluster size *k* ≥ 28 (*p*_cluster_ ≤ 0.05). (**a-c**) Contrast between confident near-threshold hits and misses with focus on (**a**) the precuneus (PCUN) and the intraparietal sulcus (IPS), (**b**) the posterior cingulate cortex (PCC; *x* = −7), and (**c**) the left anterior insula (lINS; *z* = −3). (**d-f**) Contrast between near-threshold misses and correct rejections (CR) of trials without stimulation. (**d**) Coronal view (*y* = −26) with the contralateral and ipsilateral secondary somatosensory cortices (cS2; iS2). (**e**) Sagittal view (*x* = 41) on the cS2 cluster reaching into insular cortex. (**f**) Axial view on cS2 and iS2 (*z* = 18). Left (L), right (R), the left hemisphere (LH), and the right hemisphere (RH) are indicated.

**Figure 5.**
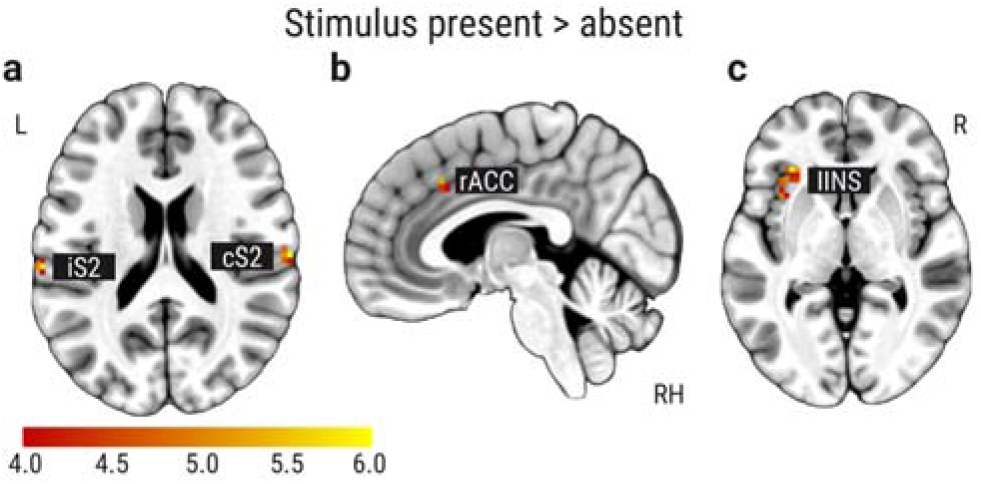
BOLD amplitude contrast for near-threshold stimulation. Correction for multiple comparison with *t*_voxel_(30) ≥ 3.92, *p*_voxel_ ≤ 0.0005 and cluster size *k* ≥ 5 (*p*_cluster_ ≤ 0.05). (**a-c**) Contrast between near-threshold stimulation trials and trials without stimulation with significant clusters in (**a**) the ipsilateral and contralateral secondary somatosensory cortex (iS2, cS2; *z* = 19), (**b**) the right anterior cingulate cortex (rACC; *x* = 4), and (**c**) the left anterior insula (lINS; *z* = 0). Left (L), right (R), and the right hemisphere (RH) are indicated.

**Figure 6.**
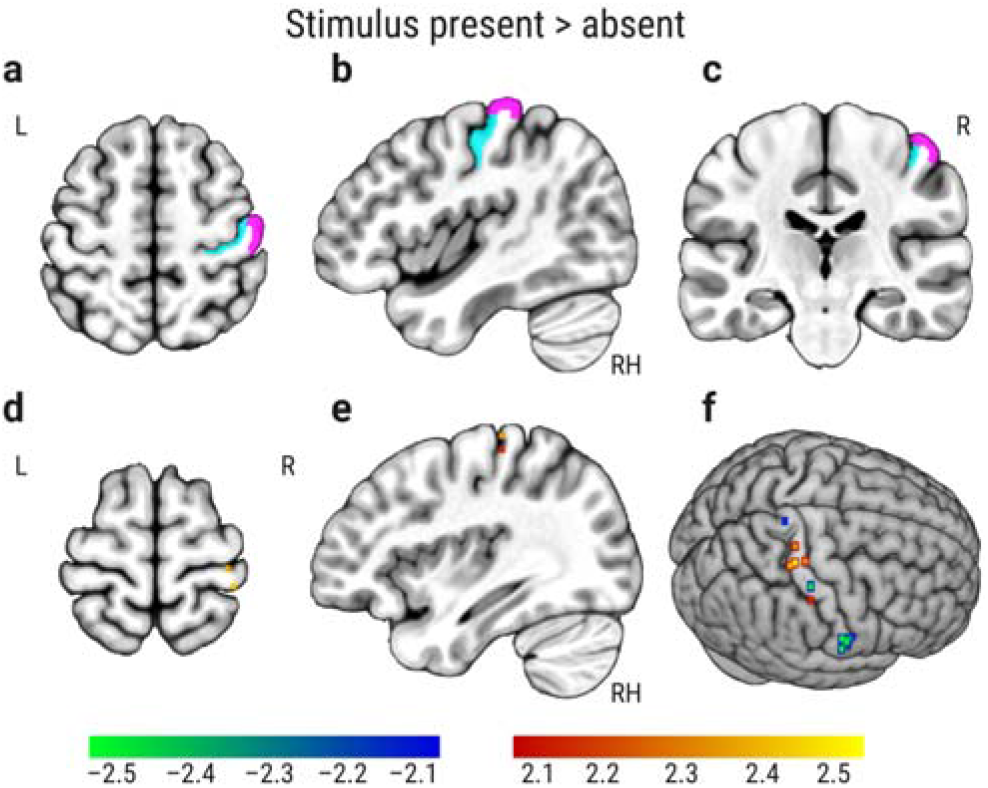
BOLD amplitude contrast for near-threshold stimulation in the primary somatosensory cortex (Area 3b and Area 1). (**a-c**) Region of interest Area 3b (cyan) and Area 1 (violet, *z* = 56, *x* = 43, *y* = −22). (**d-f**) Only voxels with *t*_voxel_(30) ≤ −2.045 and *t*_voxel_(30) ≥ 2.045, *p*_voxel_ ≤ 0.05 (*z* = 68, *x* = 36). Left (L), right (R), and the right hemisphere (RH) are indicated.

Contrasting near-threshold hits and misses (stimulus awareness) showed a fronto-parietal network including the left inferior frontal gyrus (lIFG), the left nucleus accumbens (lNAC), the left and right anterior insula (lINS1; rINS), the left and right intraparietal sulcus (lIPS1; lIPS2; rIPS) and the right precuneus (rPCUN; Figure 3a-c, Table 1). When the statistical threshold for the family-wise error was set to *p*_cluster_ ≤ 0.06, resulting in a decreased cluster size *k* ≥ 28, two additional clusters were observed for hits compared to misses in the contralateral secondary somatosensory cortex (cS2) and the left precuneus (lPCUN). When comparing missed near-threshold trials with correctly rejected null trials (somatosensory processing of undetected stimuli), the contra- and ipsilateral S2 (cS2b; iS2), the left anterior insula (lINS2) and the left supplementary motor area (lSMA) showed statistically significant activations (Figure 3d-f).

**Table 1.**
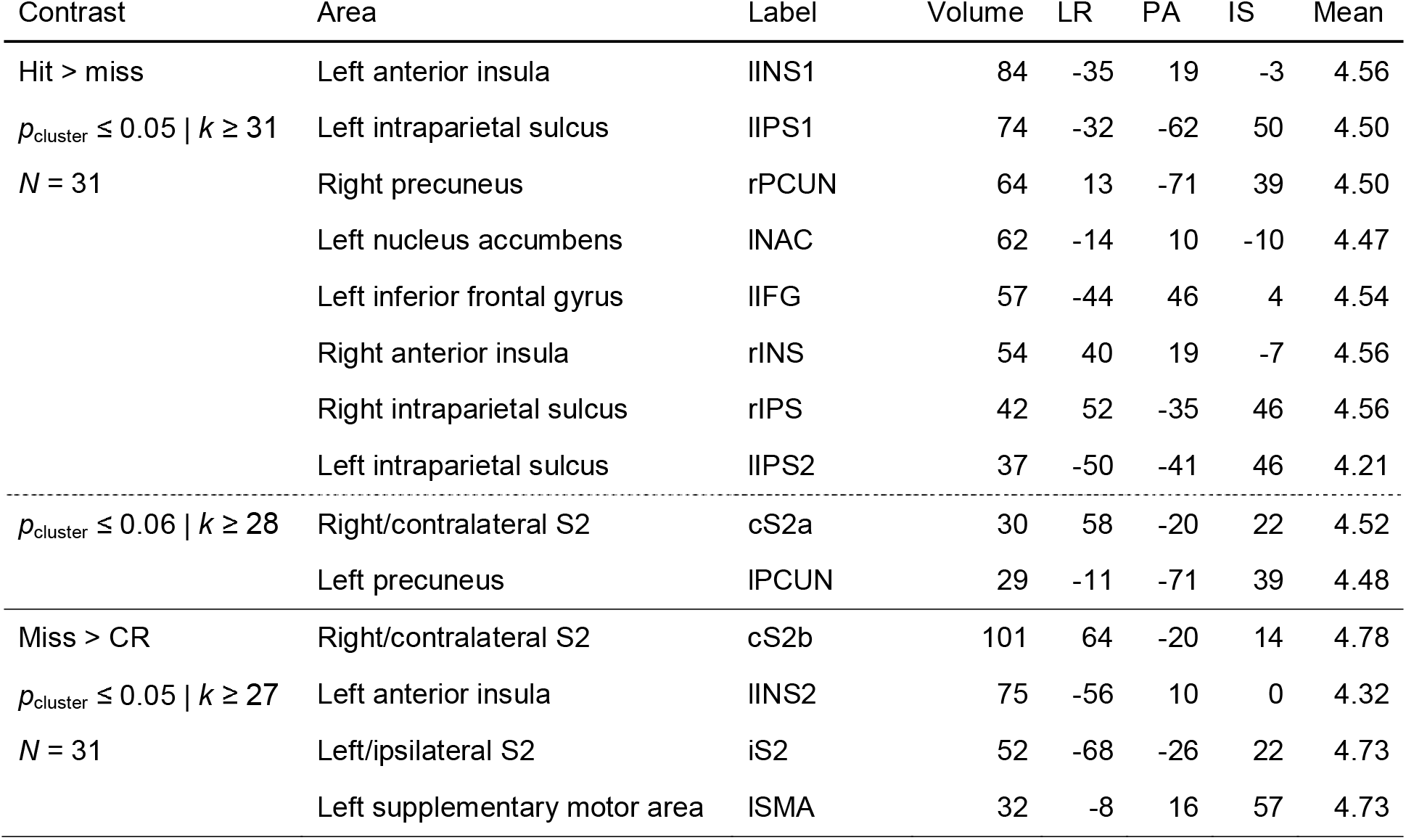
MNI coordinates for significant BOLD contrast clusters “hit > miss” and “miss > correct rejection (CR)” in Figure 3. Correction for multiple comparisons with *t*_voxel_(30) ≥ 3.92, *p*_voxel_ ≤ 0.0005 and *p*_cluster_ ≤ 0.05, resulting in a cluster size *k* ≥ 31 for “hit > miss” and a cluster size *k* ≥ 27 for “miss > CR”. Clusters are ordered by volume (number of voxels). MNI coordinates of the maximum *t* value (peak) are reported in millimeters (mm) on the left-right (LR), posterior-anterior (PA) and inferior-superior (IS) axes. The mean *t* value is the average across all voxels of one cluster.

| Contrast | Area | Label | Volume | LR | PA | IS | Mean |
| --- | --- | --- | --- | --- | --- | --- | --- |
| Hit > miss | Left anterior insula | lINS1 | 84 | -35 | 19 | -3 | 4.56 |
| $p_{\text{cluster}} \leq 0.05 \mid k \geq 31$ | Left intraparietal sulcus | lIPS1 | 74 | -32 | -62 | 50 | 4.50 |
| $N = 31$ | Right precuneus | rPCUN | 64 | 13 | -71 | 39 | 4.50 |
|  | Left nucleus accumbens | lNAC | 62 | -14 | 10 | -10 | 4.47 |
|  | Left inferior frontal gyrus | lIFG | 57 | -44 | 46 | 4 | 4.54 |
|  | Right anterior insula | rINS | 54 | 40 | 19 | -7 | 4.56 |
|  | Right intraparietal sulcus | rIPS | 42 | 52 | -35 | 46 | 4.56 |
|  | Left intraparietal sulcus | lIPS2 | 37 | -50 | -41 | 46 | 4.21 |
| $p_{\text{cluster}} \leq 0.06 \mid k \geq 28$ | Right/contralateral S2 | cS2a | 30 | 58 | -20 | 22 | 4.52 |
|  | Left precuneus | lPCUN | 29 | -11 | -71 | 39 | 4.48 |
| Miss > CR | Right/contralateral S2 | cS2b | 101 | 64 | -20 | 14 | 4.78 |
| $p_{\text{cluster}} \leq 0.05 \mid k \geq 27$ | Left anterior insula | lINS2 | 75 | -56 | 10 | 0 | 4.32 |
| $N = 31$ | Left/ipsilateral S2 | iS2 | 52 | -68 | -26 | 22 | 4.73 |
|  | Left supplementary motor area | lSMA | 32 | -8 | 16 | 57 | 4.73 |

Second, we contrasted only confident hits, misses, and correct rejections. Trials were classified as confident when rated with 3 or 4 (“rather confident” or “very confident”). Since the first trial of each block was not considered for the fMRI analysis, the participants (*N* = 31) had on average 28 confident hits (*SD* = 14), 28 confident misses (*SD* = 15), and 29 confident correct rejections (*SD* = 7). For confident hits and misses, the precuneus bilaterally (PCUN), the left and the right intraparietal sulcus (lIPS, rIPS1, rIPS2), the posterior cingulate cortex (PCC) and the left anterior insula (lINS) had significant activation clusters with conscious tactile perception (Figure 4a-c). The contralateral secondary somatosensory cortex (cS2) showed activation again with the statistical threshold *p*_cluster_ ≤ 0.06 (Table 2). Confident misses showed a higher activation than confident correct rejections in the ipsilateral and contralateral secondary somatosensory cortices (iS2, cS2). The cS2 cluster was reaching into the posterior insular cortex (Figure 4d-f).

**Table 2.** MNI coordinates for significant BOLD contrast clusters “confident hit > miss” and “confident miss > correct rejection (CR)” in Figure 4. Correction for multiple comparisons with *t*_voxel_(30) ≥ 3.92, *p*_voxel_ ≤ 0.0005 and *p*_cluster_ ≤ 0.05, resulting in a cluster size *k* ≥ 28. Clusters are ordered by volume (number of voxels). MNI coordinates of the maximum *t* value (peak) are reported in millimeters (mm) on the left-right (LR), posterior-anterior (PA) and inferior-superior (IS) axes. The mean *t* value is the average across all voxels of one cluster.

| Contrast | Area | Label | Volume | LR | PA | IS | Mean |
| --- | --- | --- | --- | --- | --- | --- | --- |
| Confident hit > miss | Left/right precuneus | PCUN | 387 | -8 | -74 | 39 | 4.66 |
| $p_{\text{cluster}} \leq 0.05 \mid k \geq 28$ | Left intraparietal sulcus | lIPS | 137 | -47 | -53 | 50 | 4.43 |
| $N = 31$ | Left anterior insula | lINS | 57 | -32 | 28 | -3 | 4.68 |
|  | Right intraparietal sulcus | rIPS1 | 42 | 55 | -38 | 50 | 4.46 |
|  | Posterior cingulate cortex | PCC | 39 | 4 | -35 | 22 | 4.45 |
|  | Right intraparietal sulcus | rIPS2 | 34 | 40 | -62 | 53 | 4.33 |
| $p_{\text{cluster}} \leq 0.06 \mid k \geq 26$ | Right/contralateral S2 | cS2 | 26 | 61 | -20 | 22 | 4.36 |
| Confident miss > CR | Right/contralateral S2 | cS2 | 141 | 64 | -20 | 14 | 4.56 |
| $p_{\text{cluster}} \leq 0.05 \mid k \geq 28$ | Left/ipsilateral S2 | iS2 | 85 | -65 | -26 | 22 | 4.60 |

Third, we contrasted all near-threshold and catch trials independent of their behavioral response to investigate the stimulation effect in the whole brain. In contrast to the BOLD contrast analysis above, the preprocessing excluded smoothing, scaling or nuisance regression to align it with the preprocessing of the functional connectivity analysis. For the near-threshold stimulation compared to no stimulation, the ipsilateral and contralateral secondary somatosensory (iS2, cS2), the left anterior insula (lINS) and the right anterior cingulate cortex (rACC) showed significant clusters with a larger activation (Figure 5, Table 3).

**Table 3.** MNI coordinates for significant BOLD contrast clusters “near-threshold stimulation > trials without stimulation (catch trials)” in Figure 5. Correction for multiple comparisons with *t*_voxel_(30) ≥ 3.92, *p*_voxel_ ≤ 0.0005 and *p*_cluster_ ≤ 0.05, resulting in a cluster size *k* ≥ 5. Clusters are ordered by volume (number of voxels). MNI coordinates of the maximum *t* value (peak) are reported in millimeters (mm) on the left-right (LR), posterior-anterior (PA) and inferior-superior (IS) axes. The mean *t* value is the average across all voxels of one cluster.

| Contrast | Area | Label | Volume | LR | PA | IS | Mean |
| --- | --- | --- | --- | --- | --- | --- | --- |
| Near > catch trials | Left anterior insula | lINS | 35 | -35 | 13 | 4 | 4.64 |
| $p_{\text{cluster}} \leq 0.05 \mid k \geq 5$ | Right anterior cingulate cortex | rACC | 15 | 4 | 22 | 39 | 4.37 |
| $N = 31$ | Right/contralateral S2 | cS2 | 13 | 64 | -20 | 18 | 5.07 |
|  | Left/ipsilateral S2 | iS2 | 6 | -65 | -26 | 18 | 4.62 |

Fourth, we contrasted all near-threshold and catch trials independent of their behavioral response only within the right primary somatosensory cortex (Area 3b and Area 1). The region of interest was defined by a multi-modal brain atlas (Glasser et al., 2016). Furthermore, the preprocessing did not include smoothing, scaling, or nuisance regression. We found positive and negative significant voxels for uncorrected *p_voxel_* ≤ 0.05 in Area 3b and Area 1. A positive voxel in the latter (*x* = 38, *y* = −38, *z* = 67; Figure 6) was close to previously reported peak coordinates in Area 1 (*x* = 38, *y* = −40, *z* = 66) for electrical stimulation of the median nerve (Schröder et al., 2019). Yet, these voxels did not meet the criteria by a correction for multiple comparisons (FDR-corrected *p* ≤ 0.05 or a cluster size *k* ≥ 12 for *p_cluster_* ≤ 0.05; Table 4). Additionally, a contrast of supra-threshold hits and correct rejections corrected for multiple comparisons in the whole brain (*p*_*voxel*_ < 0.0005 and cluster size *k* ≥ 5, *p*_*cluster*_ ≤ 0.05) revealed a cluster in the contralateral primary somatosensory cortex (*k* = 6 voxels, mean *t* = 4.27) whose peak coordinates (*x* = 52, *y* = −32, *z* = 53) were in Area 1 according to the Eickhoff-Zilles atlas (Eickhoff et al., 2005).

**Table 4.**
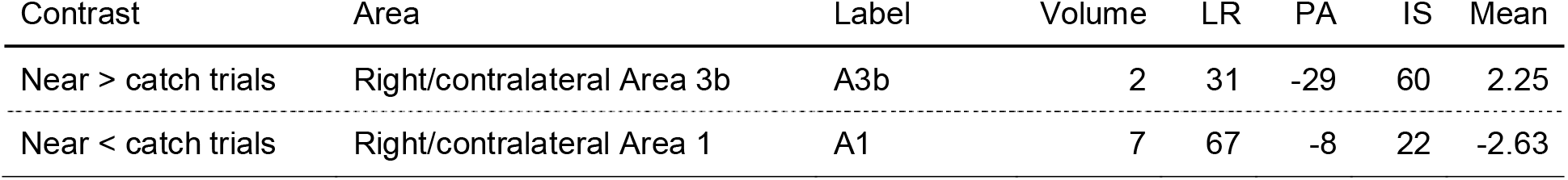
MNI coordinates for BOLD contrast clusters in the primary somatosensory cortex (Area 3b and Area 1) for “near-threshold stimulation > trials without stimulation (catch trials)” in Figure 6. Voxel threshold *t*_voxel_(30) ≥ 2.045 and *t*_voxel_(30) ≤ −2.045, *p*_voxel_ ≤ 0.05 and cluster size *k* ≥ 2. Clusters are ordered by displaying first the positive cluster and then the negative cluster. MNI coordinates of the maximum *t* value (peak) are reported in millimeters (mm) on the left-right (LR), posterior-anterior (PA) and inferior-superior (IS) axes. The mean *t* value is the average across all voxels of one cluster.

### Context-Dependent Graph Measures

We assessed whether tactile conscious perception is accompanied by alterations of the brain’s functional network topology. An atlas of 264 nodes (Power et al., 2011) was used to capture the whole-brain network as in (Godwin et al., 2015), who reported decreased modularity and increased participation with visual awareness. Whole-brain functional networks were modeled for each condition with the generalized psychophysiological interaction (gPPI; McLaren et al., 2012) without the deconvolution step (O’Reilly et al., 2012); see Methods Functional Connectivity Analysis for details). The gPPI has the advantage of controlling the context-dependent functional connectivity estimates for (a) the stimulation-related BOLD response and (b) the baseline functional connectivity across the experiment (Figure 7b). The graph-theoretical analysis of the context-dependent functional connectivity matrices was performed with the Brain Connectivity Toolbox (Rubinov and Sporns, 2010) to test for changes in the same measures of integration and segregation as in (Godwin et al., 2015). We thresholded the context-dependent connectivity matrices across a range of proportional thresholds from 5% to 40% in steps of 5% (Garrison et al., 2015) and separately calculated their normalized modularity, mean clustering coefficient, mean participation coefficient and characteristic path length (Figure 7c-f). Since Godwin et al. (2015) analyzed the graph-theoretical metrics only for confident hits and misses, we first ran the analysis for confident trials only (Figure 8). Trials were classified as confident when the yes/no-decision was rated with 3 or 4 (“rather confident” or “very confident”). Additionally, we repeated the analysis for all trials independent of their confidence rating (Figure 10) and with an extended preprocessing including a nuisance regression (Figure 11, 12).

**Figure 7.**
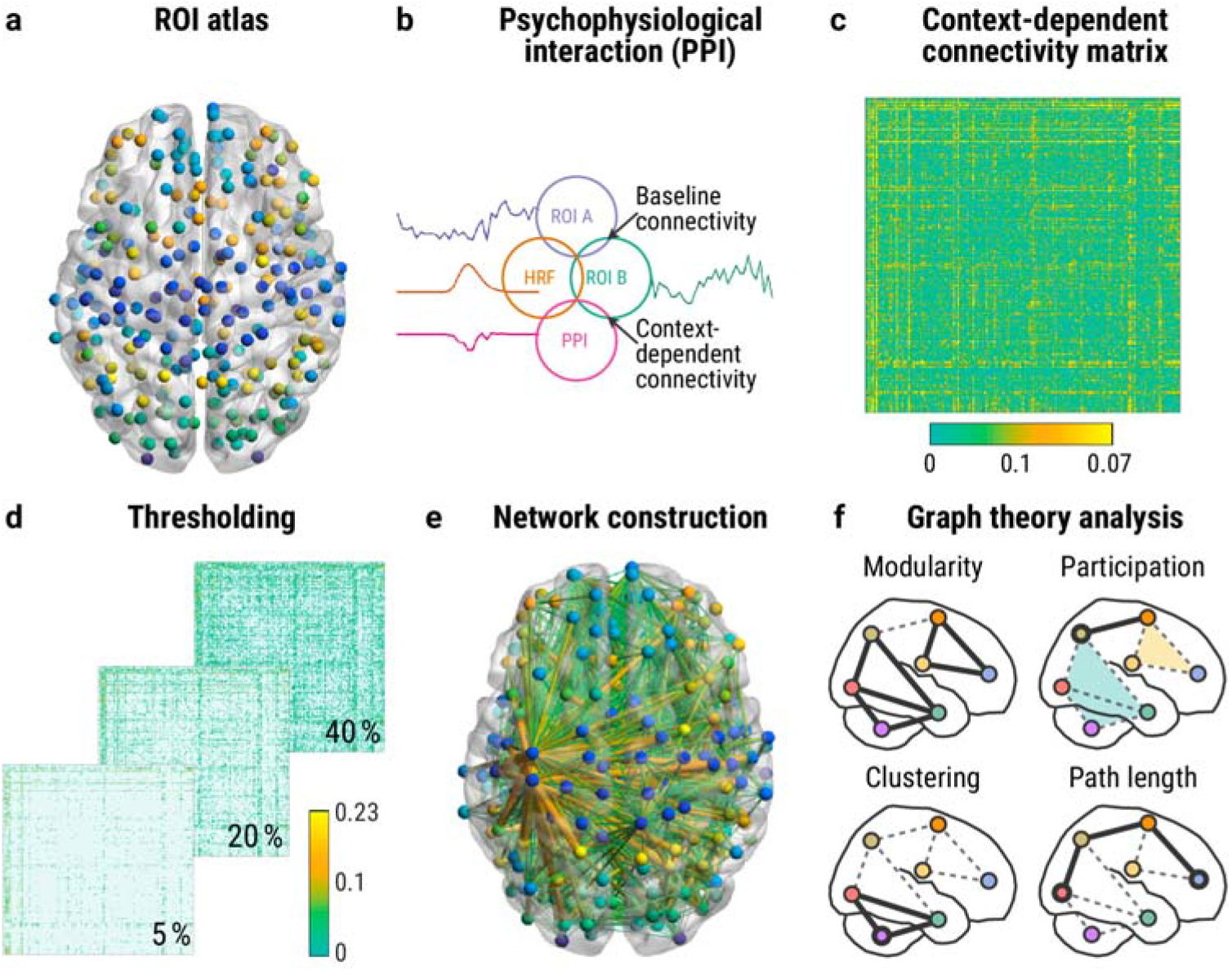
Context-dependent functional connectivity analysis. (**a**) Regions of interest (ROIs) were defined as 4-mm radius spheres at the MNI coordinates of a 264-nodes atlas (Power et al., 2011). (**b**) We used the generalized psychophysiological interaction (gPPI; McLaren et al., 2012) to calculate the context-dependent functional connectivity between all pairs of ROIs for each condition separately (hit, miss, and correct rejection). This measure controls for baseline functional connectivity and the stimulus-evoked hemodynamic response (HRF). (**c**) These context-dependent functional connectivity estimates were merged into individual, normalized, symmetric functional connectivity matrices to evaluate their network topology. For the latter, the matrices were thresholded to include only the strongest edges (**d**), and the resulting networks (**e**) were analyzed with graph-theoretical measures (**f**). For visualization, we selected the mean context-dependent connectivity matrix for hits (**c**) and thresholded it proportionally with 5-40% (**d**) and with 5% for the visualization of the edges (**e**). Edge color and diameter capture the strength of functional connectivity. Figure concept was inspired by Figure 2 in (Uehara et al., 2014).

**Figure 8.**
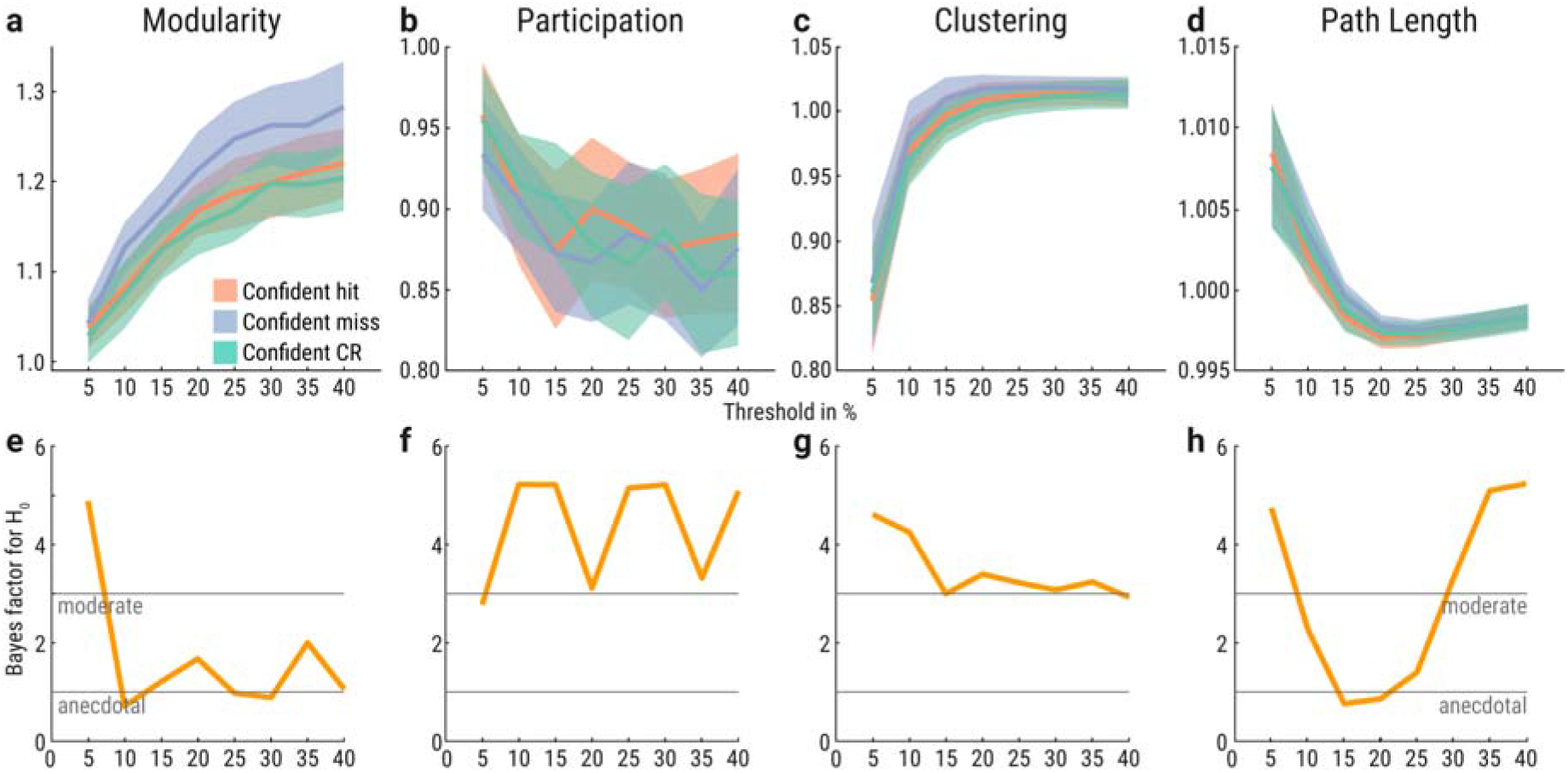
Functional network topology of only confident hits (red), misses (purple), and correct rejections (green). (**a-d**) Graph measures for network thresholds from 5-40% in 5% steps (x-axes). Y-axes indicate normalized graph metric values. Confidence bands reflect within-participant 95% confidence intervals. (**e-h**) Bayes factors (BF_01_) based on paired *t*-test between confident hits and misses. Bayes factor of 2 indicates that the evidence for the null hypothesis is twice as likely compared to the alternative hypothesis given the data. Bayes factors between 1-3 are interpreted as anecdotal and between 3-10 as moderate evidence for the null hypothesis (Schönbrodt and Wagenmakers, 2018).

Confident hits and misses showed no significant differences in measures of global segregation into distinct networks (modularity), local segregation (clustering), integration (path length), and centrality (participation) based on paired two-sided Wilcoxon’s signed-rank tests and FDR-correction (Figure 8a-d). Additionally, we calculated the Bayes factors based on paired *t*-tests with a JZS prior (*r* = √2/2; Rouder et al., 2012) to evaluate the evidence for the null hypothesis (*H*_0_: confident hits and misses do not differ). For modularity, participation, clustering, and path length, the evidence was anecdotal or moderate for the null hypothesis across the network thresholds (Figure 8e-h). The Bayes factor for modularity was below 1 at the 10%-, 25%- and 30%-threshold (Figure 8e) and hence reflecting anecdotal evidence for the alternative hypothesis. Path length had a Bayes factor below 1 at the 20%-threshold (Figure 8h). Confident correct rejections showed no significant differences to confident misses or confident hits (Figure 8a-d). Furthermore, we calculated the mean connectivity matrices of each condition for the 10%-network threshold to visualize the context-dependent functional connectivity estimates across all 264 nodes (Figure 9).

**Figure 9.**
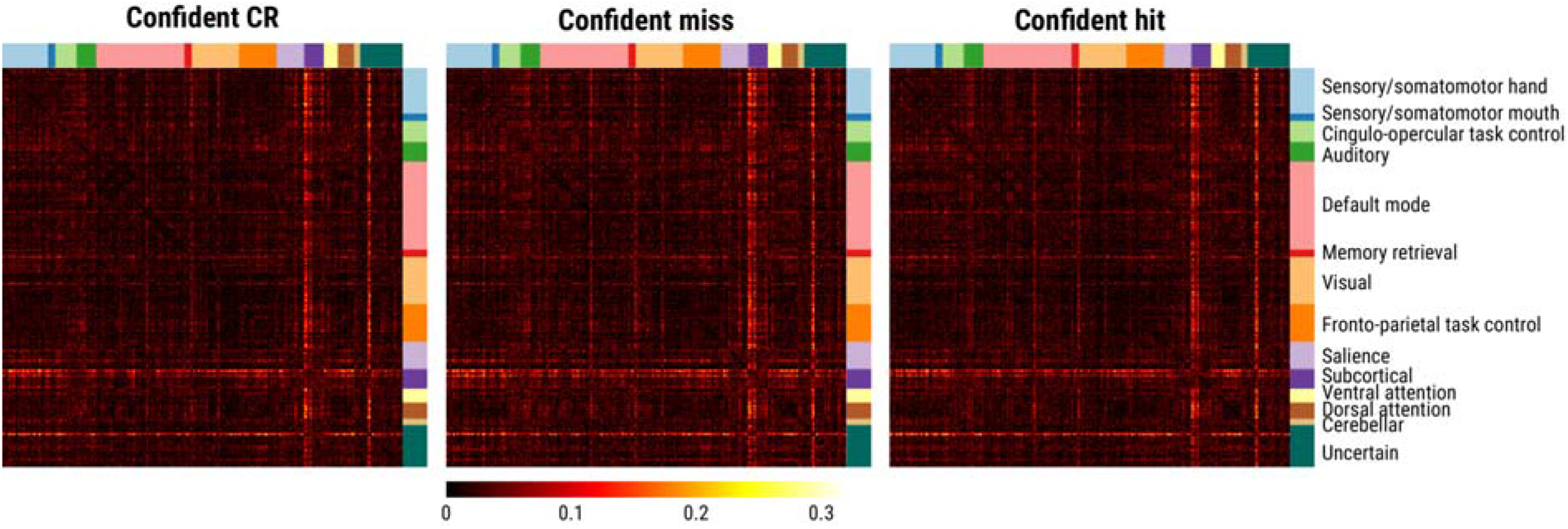
Mean connectivity matrices of confident correct rejections (CR), misses and hits for the 10%-threshold (Figure 8). The values represent the normalized gPPI estimates between the 264 nodes ordered by subnetworks.

Because graph metrics of the whole brain might be similar between conditions while the graph metrics of individual nodes and subnetworks differ, we normalized the averaged participation and clustering coefficients separately for each of the 14 Power et al. (2011) subnetworks. For this analysis, we chose the networks thresholded for the top 10% connections (Figure 9). The resulting graph metrics were compared with Wilcoxon’s signed-rank tests between the three conditions while correcting for multiple comparisons with a false discovery rate of 5%. There was no significant difference in any subnetwork between confident correct rejections, misses, and hits.

We repeated the analysis for all trials independent of the confidence rating to increase the number of trials and hence the statistical power. As in the preceding analysis (Figure 8), we observed no significant differences in modularity, participation, clustering, and path length based on paired two-sided Wilcoxon’s signed-rank tests and FDR-correction (Figure 10a-d). There was anecdotal to moderate evidence for the null hypothesis (*H*_0_: hits and misses do not differ; Fig 9e-h). Only the Bayes factors for participation at the 40%-threshold (Figure 10f) and path length at the 20%-threshold (Figure 10h) were below 1 and hence reflected anecdotal evidence for the alternative hypothesis. Correct rejections showed no significant differences to misses or hits (Figure 10a-d).

**Figure 10.**
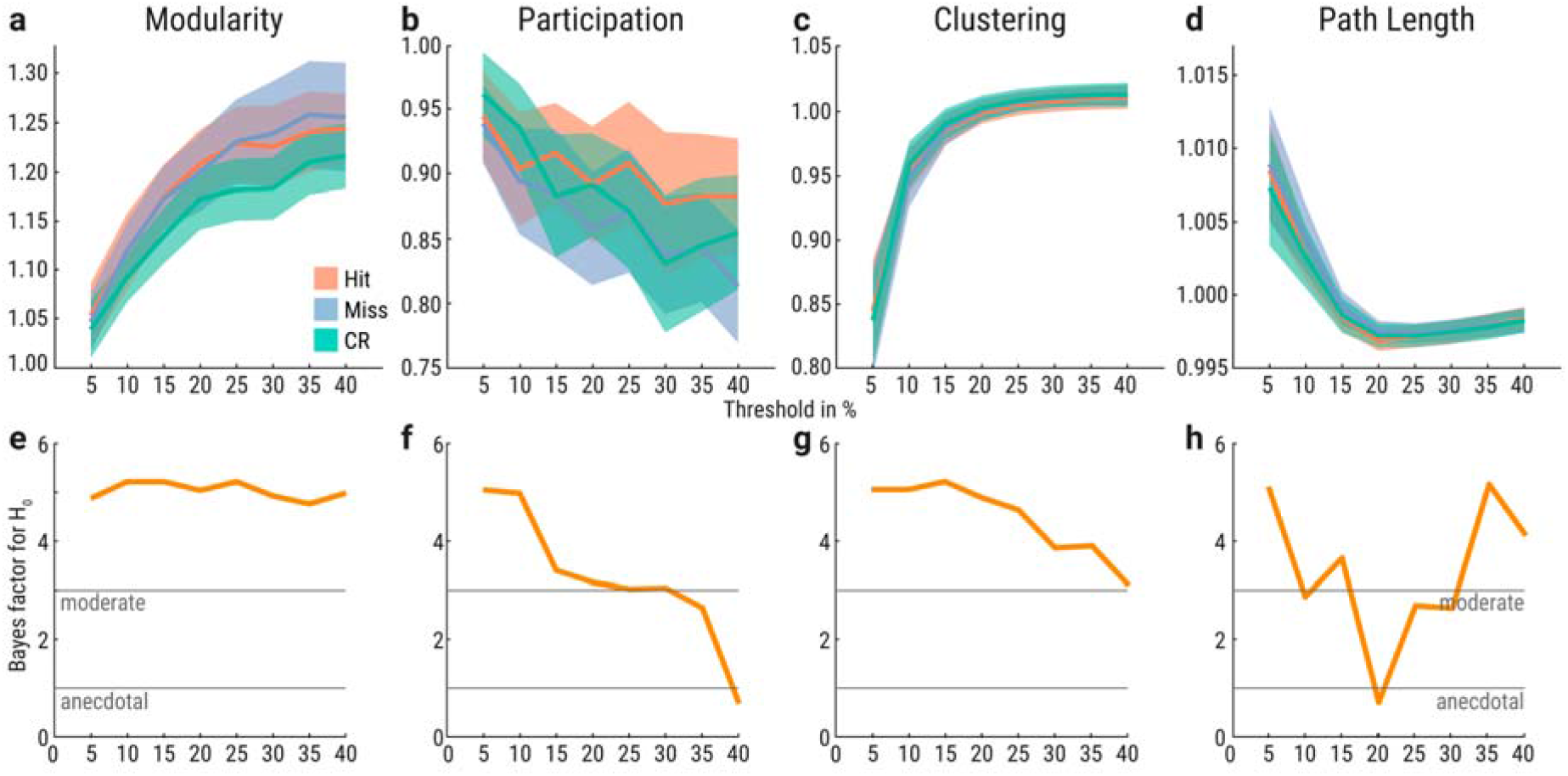
Functional network topology of all hits (red), misses (purple), and correct rejections (green). (**a-d**) Graph measures for network thresholds from 5-40% in 5% steps (x-axes). Y-axes indicate normalized graph metric values. Confidence bands reflect within-participant 95% confidence intervals. (**e-h**) Bayes factors (BF_01_) based on paired *t*-tests between detected and undetected near-threshold trials. Bayes factor of 2 indicates that the evidence for the null hypothesis is twice as likely compared to the alternative hypothesis given the data. Bayes factors between 1-3 are interpreted as anecdotal and between 3-10 as moderate evidence for the null hypothesis (Schönbrodt and Wagenmakers, 2018).

In a third step, we extended the preprocessing to include smoothing, scaling and a nuisance regression as in the whole-brain BOLD contrast analysis (Figure 3, 4). We then analyzed again the graph metrics for only confident trials (Figure 11) and all trials independent of the confidence response (Figure 12). For only confident trials, we observed no significant differences in modularity, participation, clustering, and path length based on paired two-sided Wilcoxon’s signed-rank tests and FDR-correction (Figure 11a-d). There was anecdotal to moderate evidence for the null hypothesis (*H*_0_: confident hits and misses do not differ; Figure 8e-h). Confident correct rejections showed no significant differences to confident misses or confident hits (Figure 11a-d).

**Figure 11.**
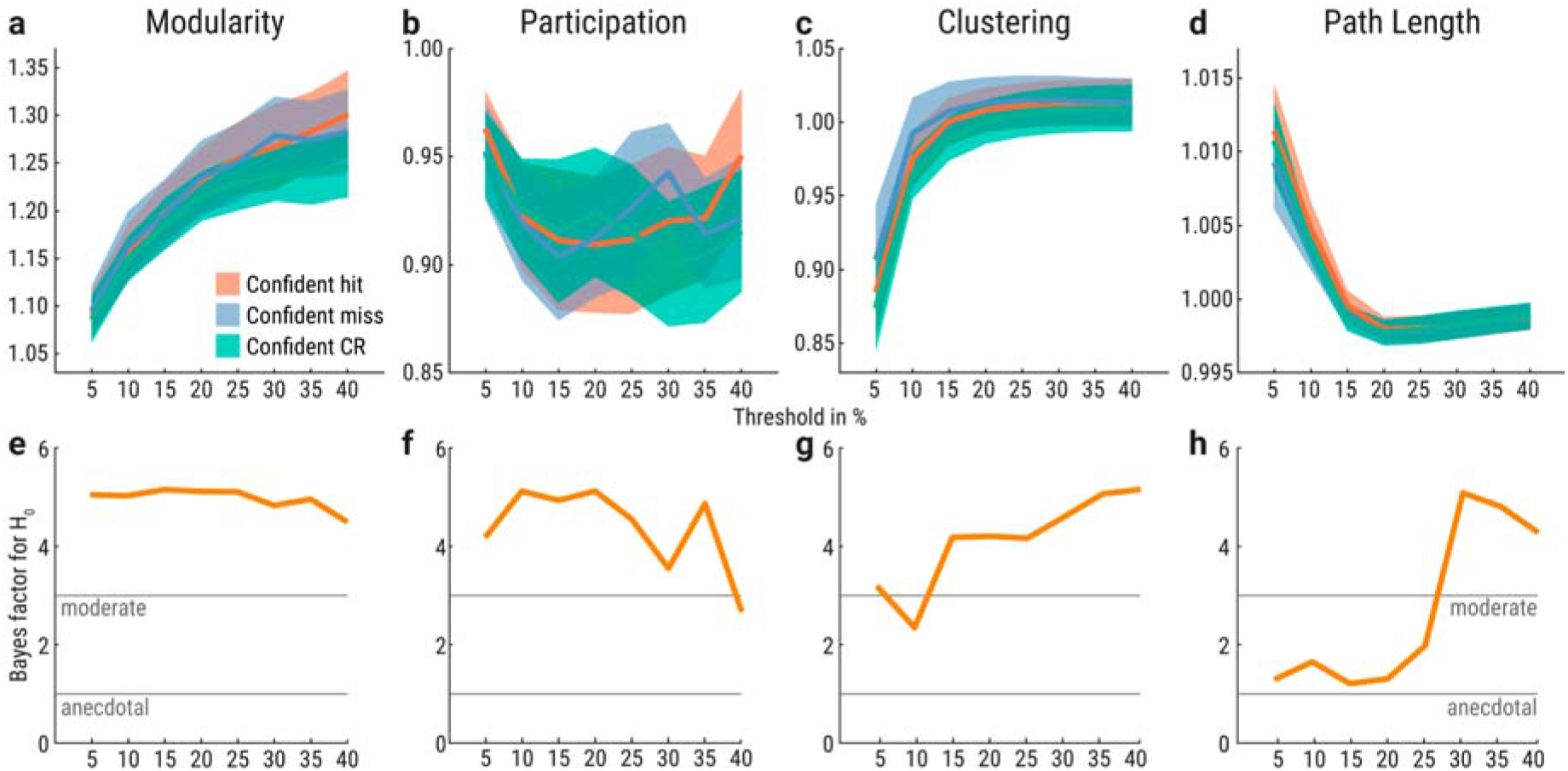
Functional network topology of only confident hits (red), misses (purple), and correct rejections (green) with extended preprocessing. (**a-d**) Graph measures for network thresholds from 5-40% in 5% steps (x-axes). Y-axes indicate normalized graph metric values. Confidence bands reflect within-participant 95% confidence intervals. (**e-h**) Bayes factors (BF_01_) based on paired *t*-tests between confident detected and undetected near-threshold trials. Bayes factor of 2 indicates that the evidence for the null hypothesis is twice as likely compared to the alternative hypothesis given the data. Bayes factors between 1-3 are interpreted as anecdotal and between 3-10 as moderate evidence for the null hypothesis (Schönbrodt and Wagenmakers, 2018).

**Figure 12.**
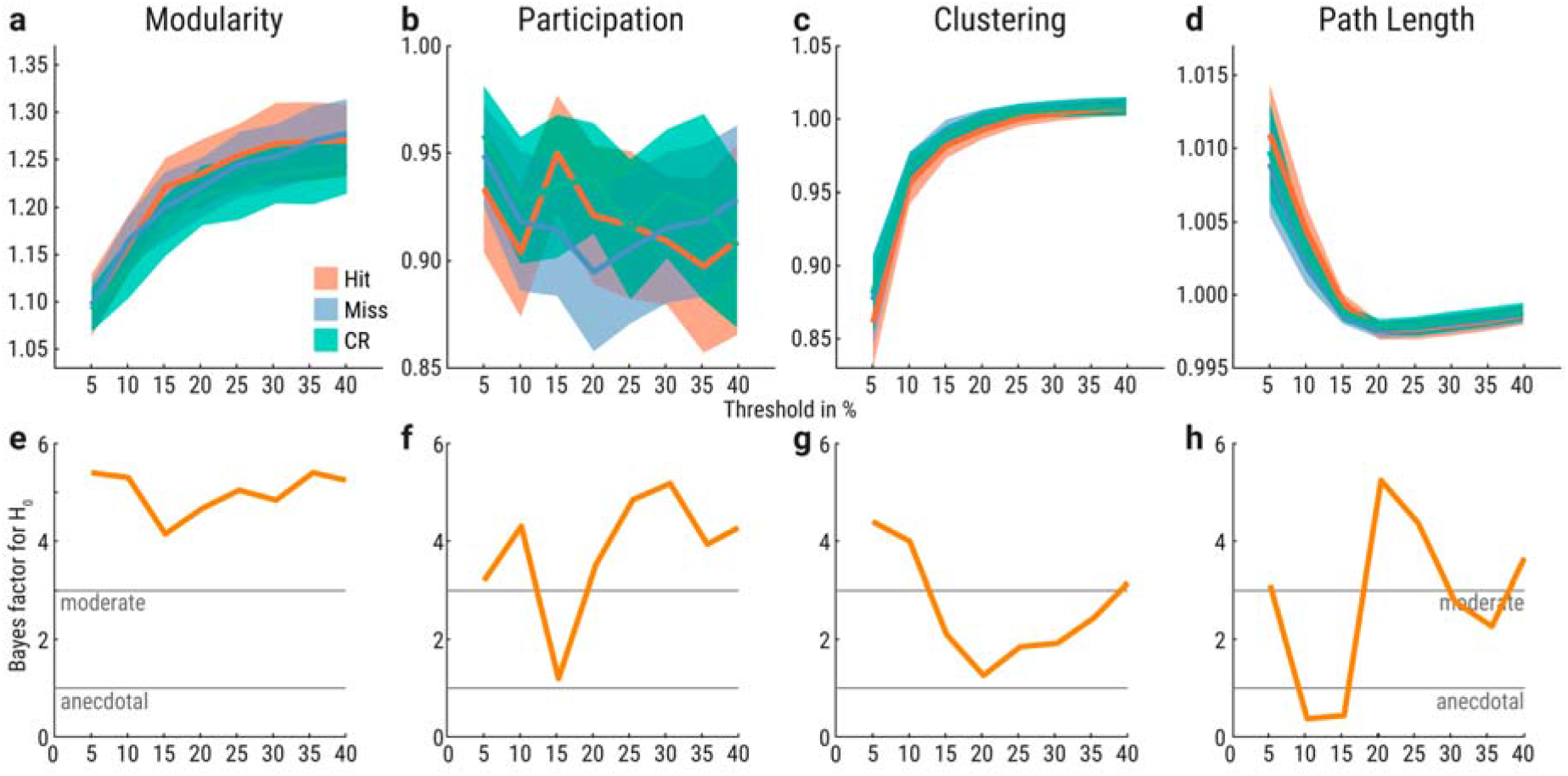
Functional network topology of all hits (red), misses (purple), and correct rejections (green) with extended preprocessing. (**a-d**) Graph measures for network thresholds from 5-40% in 5% steps (x-axes). Y-axes indicate normalized graph metric values. Confidence bands reflect within-participant 95% confidence intervals. (**e-h**) Bayes factors (BF_01_) based on paired *t*-tests between detected and undetected near-threshold trials. Bayes factor of 2 indicates that the evidence for the null hypothesis is twice as likely compared to the alternative hypothesis given the data. Bayes factors between 1-3 are interpreted as anecdotal and between 3-10 as moderate evidence for the null hypothesis (Schönbrodt and Wagenmakers, 2018).

For the extended preprocessing and all trials independent of the confidence response, hits and misses did not differ significantly in modularity, participation, clustering, and path length based on paired two-sided Wilcoxon’s signed-rank tests and FDR-correction (Figure 12a-d). There was anecdotal or moderate evidence for the null hypothesis (*H*_0_: hits and misses do not differ; Figure 12e-h). Only the Bayes factors for path length at the 10%- and 15%-threshold were below 1 (Figure 12h) and hence reflected anecdotal evidence for the alternative hypothesis. The path length was higher for correct rejections than hits at the 35%-threshold (FDR-corrected *p* = 0.017), and at the 40%-threshold (FDR-corrected *p* = 0.042).

Furthermore, to investigate whether the atlas-based approach missed functional connectivity of subnetworks, we performed a seed-based gPPI analysis with the cS2 cluster from the contrast between near-threshold and catch trials (Figure 5; *x* = 64, *y* = −20, *z* = 18). We created a 4-mm radius sphere and extracted the mean BOLD time course, as in the atlas-based approach. After computing individual gPPI models, we tested the psychophysiological interaction regressors of the three conditions (confident correct rejections, misses, and hits) against each other across participants for all voxels and applied a cluster correction for multiple comparisons (*p_voxel_* ≤ 0.0005, cluster size *k* ≥ 4 for *p_cluster_* ≤ 0.05). We did not observe any significant cluster for the cS2 functional connectivity contrast between confident hits and misses, and confident hits and correct rejections. For the contrast of confident misses and correct rejections, we found one negative significant cluster at threshold (cluster size *k* = 4 voxels) in the right cerebellum (*x* = 7, *y* = −41, *z* = 17). We repeated the analysis with extended preprocessing (including smoothing, scaling, and nuisance regression) and with all trials independent of their confidence response (corresponding to Figure 11). We did not observe any significant cluster for the cS2 functional connectivity contrast between any two of the three conditions (correct rejections, misses, and hits) for *p_voxel_* ≤ 0.0005 and *p_cluster_* ≤ 0.05 (cluster size *k* ≥ 18 for hits vs. misses, and cluster size *k* ≥ 19 for hits vs. correct rejections, and misses vs. correct rejections). That is why we conclude that the small negative cluster between confident misses and correct rejections is a spurious finding.

## Discussion

Using fMRI during a near-threshold somatosensory detection task, we investigated changes in local brain activity and functional brain network topology associated with conscious perception. We found that conscious somatosensory perception (‘detected’ compared to ‘undetected’ stimuli) led to higher activation in precuneus, intraparietal sulcus, insula, inferior frontal gyrus, and nucleus accumbens. The latter two showed higher activation only when all trials were included (confident and unconfident) but not with confident trials only. At a slightly looser statistical threshold (*p* = 0.06) bilateral secondary somatosensory cortex also showed higher activity during conscious perception. Significant positive voxels in contralateral S1 for near-threshold stimuli were only noted in an ROI-based analysis. The graph-theoretical analysis of network topology did not provide any evidence for a difference between aware and unaware trials in modularity, participation, clustering, nor path length. Finally, when comparing misses with correctly rejected catch trials, we found activation of S2, insula, and supplementary area; also, in this contrast, no changes in network properties were observed. Subsequently, we first discuss the observed BOLD activity patterns and then the absent graph metric changes in comparison to the findings in the visual system using a masking paradigm by Godwin et al. (2015).

It is generally accepted, that primary and secondary somatosensory cortices (S1, S2) are necessary for somatosensory processing leading to conscious perception (Hirvonen and Palva, 2016; Moore et al., 2013). It has long been established that lesions in S1 go along with hypoesthesia (Roland, 1987). Recently, we have shown in stroke patients, that - despite intact S1 - also lesions in S2 (along with anterior and posterior insula, putamen, and subcortical white matter connections to prefrontal structures) lead to impaired tactile conscious experience (Preusser et al., 2015). fMRI studies employing supra-threshold somatosensory stimulation - in passive or active designs - have consistently shown activation of S1 and S2 (Ruben et al., 2001). As we have previously shown, S1 and S2 are already affected by subthreshold stimuli (below the absolute detection threshold), which lead to a deactivation of these areas (Blankenburg et al., 2003). In the current study, we now show that near-threshold stimuli, which are not detected, i.e., below the response criterion (in signal detection theory terminology) for conscious detection, lead to an activation of S2 and insula (when compared to ‘correct rejections’ of catch trials). This leads to the interesting conclusion that non-detected stimuli lead to differential involvement of S1 and S2 depending on their intensity. Similarly, in our recent EEG study, we found that non-detected stimuli lead to a negativity 150 milliseconds after stimulation (N150) for principally detectable near-threshold stimuli but not for imperceptible stimuli intensities (Forschack et al., 2020). The N150 has been shown to originate in area S2 (Auksztulewicz et al., 2012). It will be an interesting issue for future studies whether an analysis based on objective detection paradigms (e.g., two-alternative forced-choice tasks, 2AFC) will further help to differentiate the meaning of these signal changes in S1 and S2, i.e., whether the “transition” from deactivation in S2 (and S1) to an activation is related to the “objective detection” in a criterion-free 2AFC task and the emergence of the N150. It should be noted that while others have shown that S1 represents the stimulus properties that get access to consciousness in interaction with S2 (Blankenburg et al., 2006; Moore et al., 2013; Rajaei et al., 2018; Schröder et al., 2019), we did not observe a strong stimulation effect in S1 for our near-threshold trials. This might be due to the weak stimulus intensity and stimulation “only” at the index finger in contrast to the commonly used median nerve stimulation.

For theories of consciousness, the difference between perceived versus non-perceived stimuli is most relevant. In our study, contralateral secondary somatosensory cortex (cS2) was found for both contrasts “hit > miss” and “confident hit > miss” when the statistical cluster threshold was set to *p* = 0.06. This finding is consistent with previous suggestions that there is additional activity in S2 with conscious detection. For example, a previous fMRI study on vibrotactile detection reported ipsilateral and contralateral S2 as the best correlate for detection success (Moore et al., 2013), and in another recent study bilateral S2 activity was best explained by a psychometric (detection) function (Schröder et al., 2019). One reason for detection-associated S2 activity might be recurrent processing (Lamme, 2006; van Gaal and Lamme, 2012). For example, in an EEG study, the detection of near-threshold electrical pulses to the finger was best explained by the recurrent processing between contralateral S1 and S2, as well as contralateral and ipsilateral S2 (Auksztulewicz et al., 2012). Another explanation might be that separate parts of the secondary somatosensory cortex might serve different functions. E.g., in a recent study, more inferior and superior parts of cS2 correlated with a binary detection function, and more posterior and anterior parts of cS2 correlated with a linear intensity function (Schröder et al., 2019).

While our data overall agree with the necessity of S1 and S2 activation for conscious perception, it is less clear whether activation of these areas is sufficient for conscious somatosensory perception: The fact that certain areas “best explain” detection in an fMRI study (Schröder et al., 2019), does not rule out that the activity of other areas is also involved in the conscious experience. Like several previous studies on conscious perception in other sensory domains (Bisenius et al., 2015; Dehaene and Changeux, 2011; Naghavi and Nyberg, 2005; Rees et al., 2002), we find fronto-parietal areas more active in the ‘detected versus missed’ contrast. Among these, the activations in left and right intraparietal sulcus, the bilateral posterior cingulate cortices, and the bilateral precuneus are consistent with the notion of a “posterior hot zone” for conscious experience as suggested by Koch and colleagues (Koch et al., 2016). Koch et al. (2016) argue that the increased activity that is seen with conscious perception in additional (e.g., frontal) brain areas is related to response preparation, and/or other task-related activations (confidence, etc.) as they have not been found activated in a no-response paradigm (Frässle et al., 2014). For example, in our study, the nine-second period between stimulation and response most likely leads to increased activity in areas involved in working memory (see, e.g., tactospatial sketchpad, Schmidt and Blankenburg, 2018). Recent literature has pointed out this interrelation: the default mode network (e.g., posterior cingulate cortex) supports a stronger global workspace configuration, which improves working memory performance (Vatansever et al., 2015) and might be beneficial for conscious perception. On the other hand, even if one assumes that a “pure sensory conscious experience” could arise from a “posterior hot zone” only, the increased activity in other brain areas with conscious perception - still leaves the possibility of conscious experience related to action, confidence, working memory etc. to be dependent on e.g., frontal brain areas (Frith, 2019). In this view, the “integrated conscious experience” during a task would then be related to the entire fronto-parietal network. Obviously, this notion is close to the global workspace theory (Dehaene et al., 2006; Dehaene and Changeux, 2011; Mashour et al., 2020). While our data do not allow to definitely differentiate between these major theories of consciousness, we provide new information for somatosensory conscious perception, which is consistent with the idea that domain-general areas (interacting with domain-specific areas for example via recurrent processing (Lamme, 2006; van Gaal and Lamme, 2012) play a role for conscious perception.

A major hypothesis underlying our study was that conscious perception goes along with widespread changes in graph metrics as it has recently been reported by Godwin et al. (2015) for the visual system. However, we did not observe such context-dependent functional connectivity changes that result in network topology alterations through modularity, participation, clustering and path length between hits, misses or correct rejections. This does not change when the number of trials is increased by considering all trials independent of their confidence response (Figure 10). Also, improving the preprocessing with a nuisance regression that controls for motion and noise components derived from white matter and ventricles, does not affect this result (Figure 11, 12). The isolated network topology differences between correct rejections and hits at the 35-40%-threshold for path length with the improved preprocessing (Figure 12d) were not consistent across thresholds and not present in the analysis of only confident trials (Figure 11), as well as in the analyses with the basic preprocessing (Figure 8, 10). That is why we do not interpret these differences as a valid and reliable effect. Thus, there was neither a functional network alteration by stimulus awareness (hit > miss) nor by the detected (hit > CR) or undetected stimulation (miss > CR).

Two apparent differences between the study of Godwin et al. (2015) and our study are the somatosensory versus visual modality and the use of a masking paradigm (Godwin et al.) as opposed to near-threshold stimuli. However, assuming that the connectivity changes observed by Godwin et al. are related to the conscious experience, it is difficult to see why those differences should explain the different results. One possible reason why Godwin et al. observed whole-brain network topology alterations for visual awareness might be the unbalanced physical similarity between aware and unaware trials. Hits and misses originated from two different masking conditions: backward masking generated 83% of all hits and forward masking 84% of all misses. Additionally, their total number of trials for 24 participants was not balanced (276 confident misses vs. 486 confident hits). In contrast, our study did not rely on masking the target stimulus and resulted in a balanced total amount of 882 confident misses and 870 confident hits for 31 participants (Figure 8, 10). Furthermore, we also present the results of 1507 hits, and 1190 misses independent of the confidence rating (Figure 10, 11). Future studies investigating visual awareness may be able to distill conscious percepts for present stimuli without confounding masking conditions, for instance taking advantage of sub-millisecond precision of modern tachistoscopes (Sperdin et al., 2013). Awaiting such studies as well as further studies on the somatosensory system and acknowledging – of course – that “absence of proof” is not “proof of absence” – our study cannot provide support for changes in graph metrics with awareness.

In summary (integrating our previous findings on the effect of subthreshold stimulation (Blankenburg et al., 2003; Taskin et al., 2008), there seem to be three discernable stages of fMRI-BOLD signal changes with increasing somatosensory stimulus intensity: (i) A deactivation of S1, S2 following (trains of) subthreshold (never-detected) stimuli, (ii) activation of S1, S2, and insula following near-threshold not-detected stimuli, (iii) additional activation of S1, S2 accompanied by activation of a fronto-parietal (likely to be domain-general) network when stimuli are consciously perceived. The potentially differential contribution of the involved brain areas to the conscious experience should be subject to future investigations in which modulations of different aspects of the tasks (e.g., varying delay, way of report, design e.g., 2AFC versus yes/no task) may be employed. Our study could not confirm changes in graph metrics with awareness for the somatosensory system. Whether this is related to the specific somatosensory modality (electrical nerve stimulation) or the weak stimulation should be investigated by future studies. We think that the data of electrical finger nerve stimulation - despite its limited spatial extent - is a useful model for a range of somatosensory receptors in the fingers, because the nerve integrating the receptor signals is directly stimulated. This view is also supported by similar topographical activation patterns for passive proprioceptive stimulation compared with tactile stimulation (Nasrallah et al., 2019). The potentially differential contribution of the involved brain areas to the conscious experience of electrical stimuli is in line with global broadcasting of individual content of consciousness across the brain without substantial reconfiguration of the brain's network topology resulting in an integrative conscious experience - at least for the observed somatosensory submodality.

## Acknowledgements

The research was funded by the Max Planck Society. We thank Ramona Menger, Anke Kummer, Mandy Jochemko, and Nicole Pampus for their data acquisition support; Bettina Johst, Hendrik Grunert, and Jöran Lepsien for their technical advice; Heike Schmidt-Duderstedt and Kerstin Flake for preparing the figures for publication; and Joshua Grant for proofreading and commenting.

